# Multiple elements of addiction vulnerability are independently heritable in rats

**DOI:** 10.1101/566018

**Authors:** Maya Eid, Dominika Pullmann, Hao Li, Alen Thomas, Thomas C. Jhou

## Abstract

**Background:** Although cocaine is powerfully rewarding, only a fraction of drug-exposed individuals actually transition to become drug-dependent users, possibly due in part to genetic influences, as addictions are among the most heritable of human neuropsychiatric disorders. Consistent with addiction being a complex disorder, prior studies have noted many factors that predict addiction vulnerability, such as drug-induced aversive effects, behavioral responses to novelty, and sensitivity to punishment.

**Methods:** We tested different inbred strains of rats, as well as selectively bred Sprague-Dawley (SD) or Heterogeneous Stock (HS) rats, on cocaine avoidance, sensitivity to punishment, and locomotor responses to novelty, and calculated heritability estimates of these behaviors. We tested animals on additional control tasks (progressive ratio, shock avoidance) to control for alternate interpretations of addiction-like behaviors.

**Results:** Conditioned avoidance to cocaine varied greatly between individual rats as measured using either the runway operant task or conditioned place aversion (CPA). Cocaine avoidance responded rapidly to selective breeding, yielding heritability estimates of 0.70 and 0.58 in SD and HS rats. Resistance to punishment was also highly heritable in inbred rats (estimated h^2^ = 0.62), and varied independently of cocaine avoidance despite both behaviors being mediated by the RMTg. Furthermore, cocaine avoidance correlated positively with cocaine-induced c-Fos induction in the RMTg, negatively with initial rates of acquisition of intravenous cocaine self-administration, and positively with both cued and cocaine-primed reinstatement.

**Conclusions:** Cocaine avoidance and resistance to punishment are strongly and independently heritable behaviors that may both control different aspects of individual propensity to acquire and maintain drug-seeking.

## INTRODUCTION

Over 90% of Americans have had some exposure to drugs of abuse, but only 15-32% of individuals exposed to the major classes of abused drugs go on to become dependent, with the rest presumably being able to curtail usage on their own [1-3]. The reasons for these individual differences are complex, but genetic factors likely play strong roles, as addictions are among the most heritable of human neuropsychiatric disorders, with cocaine addiction having particularly high heritability estimates (h^2^ = 0.72) [4-6]. However, the specific genes involved in human addiction have remained elusive, due in part to the complexity of addictive behaviors themselves, which likely involve multiple distinct aspects of vulnerability. For example, prior work has identified numerous factors, including response to novelty, drug-induced aversive effects, and sensitivity to punishment, that vary between individuals, are heritable, and predict drug-seeking [7]. Among these factors, cocaine’s aversive effects are somewhat understudied, relative to its rewarding effects, but are experimentally robust, with multiple studies showing that single doses of cocaine produce an initial rewarding phase followed by an aversive “crash” about 15 minutes later, that is sufficient to slow or stop cocaine-seeking in many animals [8-13]. This aversive effect depends critically on the rostromedial tegmental nucleus (RMTg), a major GABAergic midbrain input to midbrain dopamine (DA) neurons [11, 14-17], and hence we hypothesize that this initial crash may govern some of the individual variation in early acquisition of drug-seeking, and that this variation may be dependent on RMTg function.

In addition to factors regulating early acquisition of drug-seeking, addiction propensity also depends on factors affecting later use, such whether individuals curtail use as external costs of drug use increase [18]. Indeed, continued drug use despite negative consequences (financial, social, or medical) is a defining feature of human addiction [19]. In rodents, we model these effects by examining “resistance to punishment”, using a task that measures the propensity to continue to seek rewards despite highly aversive outcomes [20].

Given the evidence for heritability of addiction propensity in humans, we tested addiction heritability in rodent models that allow the dissection of addiction vulnerability into multiple constituent components, which is difficult or impossible in human populations. We particularly examined addiction vulnerability governed by cocaine’s aversive effects and by punishment learning, using Sprague Dawley rats (SD), several inbred rat strains, and heterogeneous stock (HS) rats. We found that both behaviors are highly heritable and inherit independently of each other, and that cocaine avoidance also correlated with cocaine-induced c-Fos levels in the RMTg. Finally, we noted that cocaine’s aversive effects have opposite influences on two distinct stages of drug-seeking behavior, where it seems to slow acquisition of lever pressing for cocaine, but also promotes reinstatement.

## RESULTS

### Individual Sprague-Dawley rats show large variation in cocaine-avoidance, independently of locomotor variations

We first tested 26 male and 22 female Sprague-Dawley (SD) rats on a runway operant task previously shown to be highly sensitive to the aversive effects of cocaine [8, 10], in which animals traverse a corridor to receive a single dose (0.75mg/kg) of cocaine. Consistent with prior reports, the average latency to reach the goal compartment was low for trials 1-3 and significantly higher for trials 4-7, while showing large individual variations in the degree of this increase. Across all trials, these latencies were not different between male and female rats (n=22-26 per group, t(46)=0.7035, p=0.49) **(figure 1A).** Run latencies were however higher in both males and females compared to a group of animals running to receive saline (two-way ANOVA, Bonferroni test, n=6-26 per group, p<0.01) **(figure 1B),** indicating that the increased latencies are not due to simple habituation or other non-specific reductions in exploratory activity. Because the runway apparatus is initially novel to the animal, and because novelty-induced locomotion has been shown to correlate with cocaine seeking and taking [7], all animals had also been habituated to the runway for three trials prior to the first cocaine trial to reduce novelty-related effects (see methods). To further rule out the possibility that animals with high latencies were simply less exploratory in novel environments, we explicitly measured locomotor activity in a novel operant chamber distinct from the runway apparatus and found no correlation between locomotor activity and run latencies, for either male or female rats (males: n=25, r^2^=0.06, p=0.25, females: n=18, r^2^=0.09, p=0.0.23) **(figure 1C)**.

**Figure 1.**
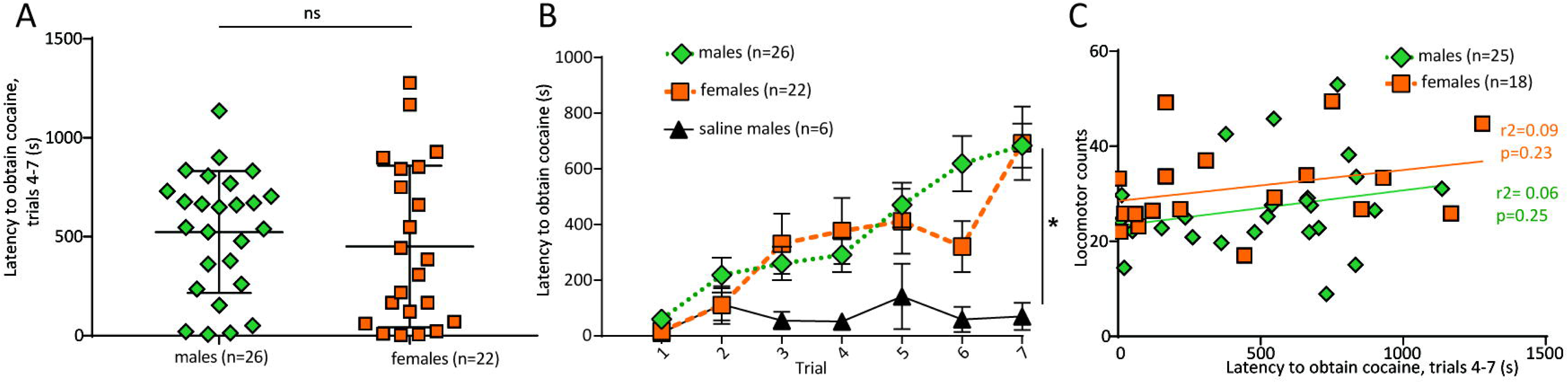
(A) Individual outbred Sprague-Dawley (SD) rats showed large variation in latency to obtain cocaine after 4-7 conditioning trials. (B) In contrast, animals receiving saline instead of cocaine showed consistently short latencies across all trials. (C) Latency to obtain cocaine did not correlate with novelty-induced locomotion, for either male or female rats. *\*p<0.05*.

### Cocaine-avoidance responds strongly to selection in the outbred SD rats

To examine whether individual differences in cocaine avoidance on the runway were heritable, we first performed selective breeding on a group of SD rats. From the 48 animals screened on the runway above, we selected two pairs each of animals in the top or bottom quartiles of the distribution of run latencies, while keeping novelty-induced locomotion scores within the middle quartiles of their distributions. This selection process was then repeated on the resultant offspring, yielding a second generation of offspring. After one generation, we observed a 3-fold difference in run latencies between offspring of high- and low-avoider parents (n=54, t(50)=3.525, p<0.0001) (males and females combined), yielding a heritability estimate of h^2^=0.70. A second generation of selective breeding produced an even larger divergence, as the average latency of high-avoider offspring was almost 14 times that of low-avoider offspring (n=32, t(31)= 6.448, p<0.0001)) **(figure 2A).** Although for the females, in contrast to the males, the divergence between high- and low-avoiders took two generations to become statistically significant, these results taken together suggest that cocaine avoidance strongly responds to selective pressure **(Figure 2B)**. As we had chosen parents that were close to the group means in novelty-induced locomotion, neither generation of offspring of high- and low-avoiders differed from each other in this parameter (one-way ANOVA, Bonferroni test, n=10-15 per group, p>0.5) **(figure 2C)**, confirming that we did not inadvertently alter this trait when selecting for cocaine avoidance. Finally, to determine if individual differences in avoidance behavior are predictive of self-administration, we attempted to train the second generation of high-avoiders on this task. Out of the 7 high-avoiders from the second generation, 5 failed to acquire lever-pressing despite 3-4 weeks of training that included standard strategies used to increase self-administration such as lever extenders or food baiting of levers [21]. Notably, food bait (combined with food deprivation) increased lever pressing on the active lever only while the bait was present, with lever pressing returning to baseline immediately after bait removal (two-way ANOVA, Bonferroni test, n=5, active lever cocaine vs cocaine+food deprivation+bait: p=0.009, cocaine+food deprivation+bait active vs inactive lever: p<0.0001) **(Figure 3D)**.

**Figure 2.**
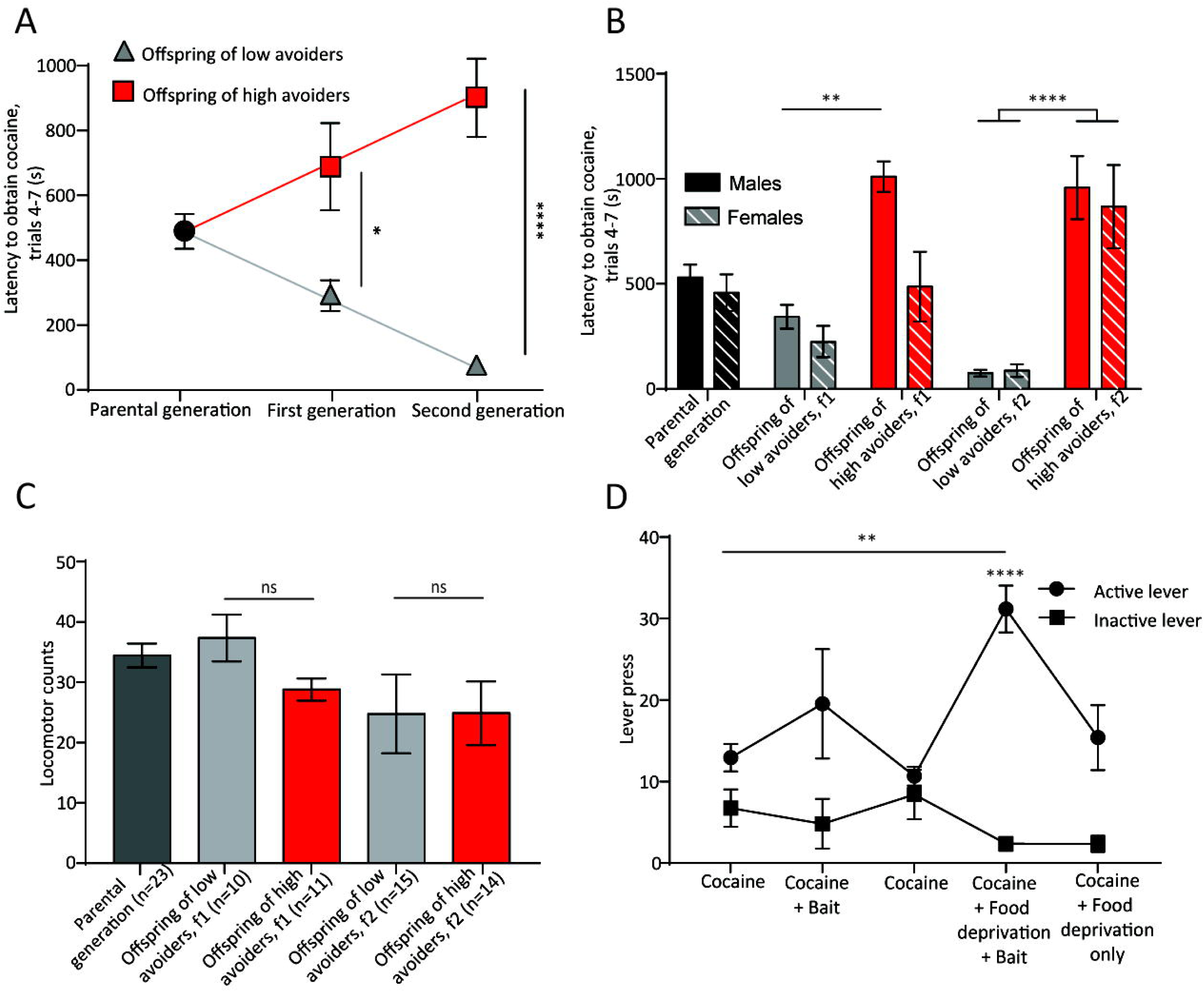
(A) SD rats responded rapidly to selection pressure for cocaine avoidance, showing large differences after both first and second generations. (B) Offspring of high and low avoiders of the second generation differed significantly in run latencies, for both male and female rats, and in the first generation for male rats only (C) Offspring of high and low avoiders did not differ in locomotor responses to novelty in either generation. (D) Offspring of high-avoiders failed to acquire lever-pressing despite training that included use of lever extenders and food baiting. *\*p<0.05, **p<0.01, ****p<0.0001.*

**Figure 3.**
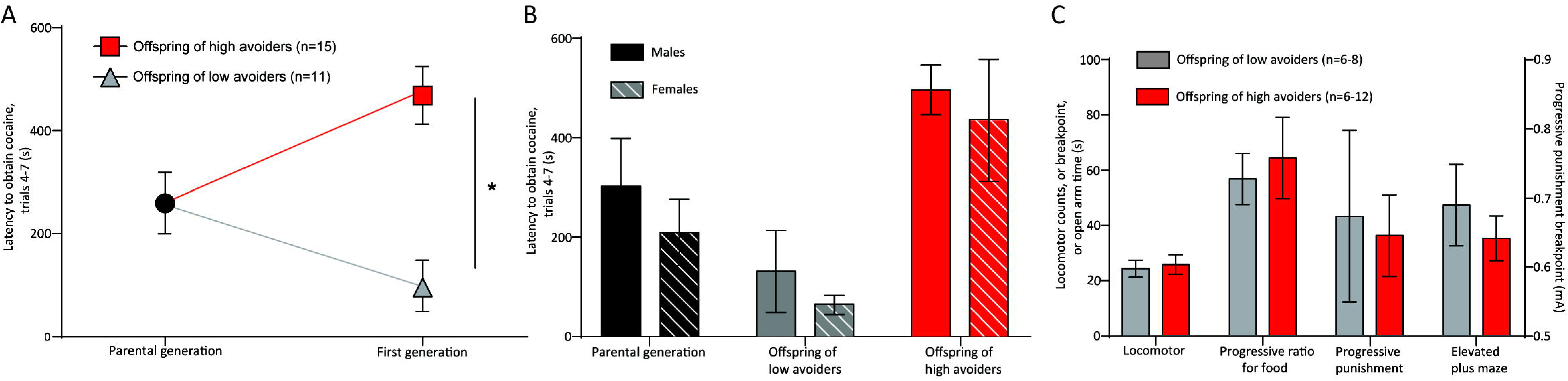
(A) HS rats responded rapidly to selection pressure for cocaine avoidance, showing large differences after only one generation (male and female offspring combined) (B) HS males and females did not differ in their latency to obtain cocaine. (C) Offspring of high- and low-cocaine avoiders were not different in progressive ratio, punishment resistance, locomotor response to novelty nor open arm time. *\*p<0.05.*

### Cocaine-avoidance responds strongly to selection in Heterogeneous Stock (HS) rats and inherits independently of control behaviors

Although we have shown evidence that cocaine avoidance is heritable in SD rats, this strain is not as well suited for fine mapping of chromosomal loci as NIH Heterogeneous Stock rats (HS), an intercross of 8 distinct inbred strains specifically designed to facilitate the fine mapping of chromosomal loci [22, 23] (A. Palmer unpublished observations). We screened 16 HS rats (8 male, 8 female) on the runway task for cocaine avoidance and generated two mating pairs each of the highest and lowest avoiding parents (4 total litters). Replicating the results from the SD breeding process, we observed that the offspring of high-avoiders had five-fold longer run latencies to obtain cocaine than the offspring of low-avoider parents (n=26, t(24)=4.909, p<0.0001) **(figure 3A)**, yielding a heritability estimate of h^2^ = 0.58. Since male and female HS did not show significant differences in run latencies in either the parental or offspring generation (two-way ANOVA, Bonferroni test, n=5-9 per group, p>0.05) **(figure 3B),** we combined both genders of offspring in all subsequent analyses. As with the SD rats, the parents were selected to be close to the median on novelty-induced locomotion. Indeed, we found no difference in novelty-induced locomotion between offspring of high- and low-avoider parents (n=8-9 per group, t(15)=0.3191, p=0.750). To assess whether differences in cocaine-seeking could have arisen from differences in generalized motivation to seek rewards, we tested animals on a progressive ratio food-seeking task, in which animals lever press for food pellets, with the required number of presses per pellet increasing progressively throughout the task. We found that offspring of high and low avoiders had similar breakpoints (n=6 per group, t(10)=0.4404, p=0.67). To examine whether differences in cocaine avoidance could have arisen from more general differences in punishment learning, we tested the animals on a *progressive punishment* task in which rats pressing for food pellets also receive concurrent footshocks whose intensity progressively increases throughout the task. in this task, a “shock breakpoint” is reached when lever pressing is largely inhibited, thus exceeding the highest amount of shock an animal will endure for a given reward. Again, this breakpoint was not different between the two offspring groups (n=6 per group, t(10)=0.4404, p=0.67). Finally, we test animals on an elevated plus maze, and found that high- and low-avoiders had no differences in anxiety levels measured by time spent in the open arms of the maze (n=7 per group, t(12)=0.2031, p=0.84) **(figure 3C)**. Altogether, these findings once again demonstrate using a different strain that cocaine avoidance is heritable, and that the differences observed in run latencies between the two offspring groups were not likely due to general differences in exploration, motivation, learning capacity or anxiety.

### Cocaine has dual rewarding and aversive effects, with the aversive component driving individual variation in drug-seeking behavior via RMTg activation

Although we had hypothesized that high-avoider rats exhibit stronger aversive effects of cocaine, their long latencies to reach the goal compartment would also be consistent with them experiencing lower rewarding effects of the drug. To address this issue, we tested the HS offspring in a conditioned place preference (CPP) task in which rats were either placed into conditioning chambers immediately after receiving an infusion of i.v. cocaine (0.75 mg/kg), or 15 minutes afterwards. We found that rats with high (> 300 s) or low (<300 s) run latencies both showed robust cocaine CPP when placed into chambers immediately after cocaine infusions. However, high-latency rats showed dramatically higher conditioned place aversion (CPA) than the low-latency rats (one-way ANOVA, Bonferroni test, n=7-8 per group, p>0.9999 and, p=0.04, respectively) **(figure 4A)**. Correlational analysis of individual animals showed similar results, in which run latencies correlated with CPA but not CPP scores (n=15, r^2^=0.48, p=0.004, r^2^=0.028, p =0.5 respectively) **(figure 4B)**. These results are consistent with prior reports [9] and suggest that individual variation in cocaine-seeking likely depends more on variation in cocaine’s aversive effects, rather than its rewarding effects, which were relatively constant across animals. Since the rostromedial tegmental nucleus (RMTg) has been previously shown to critically mediate delayed aversive responses to cocaine, we examined whether runway latencies correlated with levels of cocaine-induced RMTg c-Fos, an immediate-early gene whose expression has been used as a marker of cell activation. Indeed, RMTg c-Fos counts showed positive significant correlations with run latencies, in four independent groups of animals (vendor-purchased HS rats, vendor-purchased SD rats, offspring of high- and low-avoider HS rats, and inbred rat strains), suggesting that the higher aversive responses could be mediated by higher RMTg activity. (n=5-16 per group, p<0.05) **(figure 4C)**.

**Figure 4.**
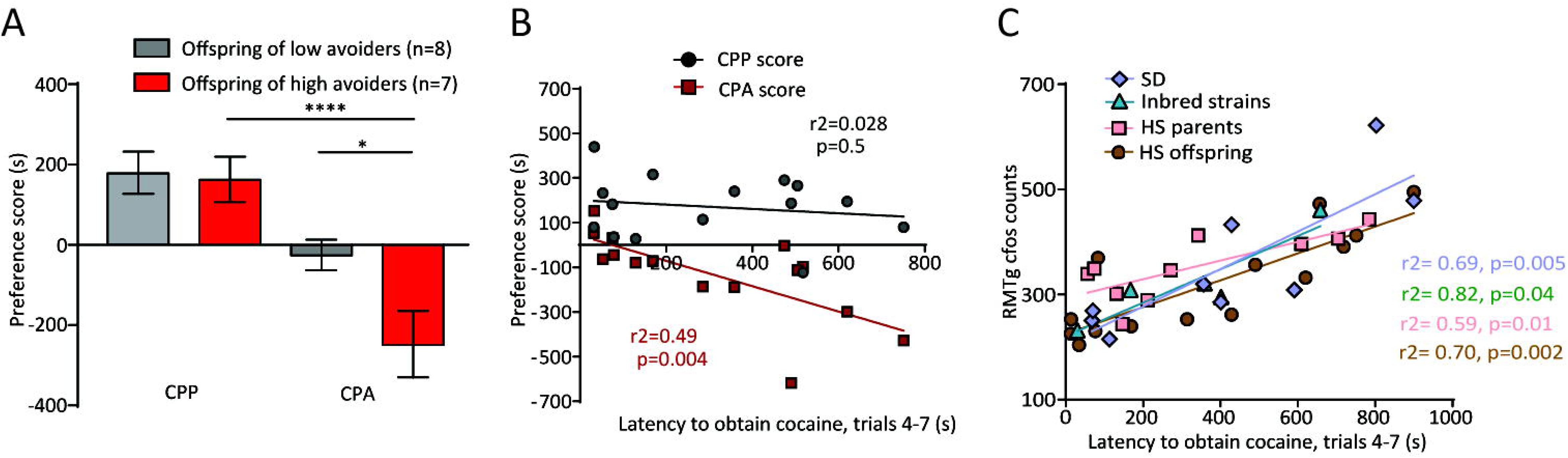
(**A**) Cocaine produces strong conditioned place preference (CPP, grey bars) for both high and low avoiders if rats are placed into conditioning chambers 0-15 minutes after iv cocaine infusions. However, high cocaine avoiders (but not low-avoiders) showed a conditioned place aversion (CPA, red bars) when rats were placed into chambers 15-30 minutes post-infusion. (**B**) Runway latencies did not correlate with CPP scores on the 0-15 minute condition, but strongly correlated with CPA scores in the 15-30 minute conditions. (C) RMTg c-Fos counts correlates with the animals’ latency to obtain cocaine in 4 independent groups: vendor-purchased HS rats, vendor-purchased SD rats, offspring of high- and low-avoider HS rats, and inbred rat strains. *\*p<0.05, ****p<0.0001.*

### Cocaine avoidance differs markedly between inbred rat strains and is not due to generalized motivational or learning differences

Because inbred rat strains have homogeneous genotypes, they can be used to separate environmental from genetic determinants of behavior, thereby providing an alternate measure of heritability independently of the selective breeding experiment. Hence, we examined 5 of the 8 constituent inbred strains comprising the HS strain: the ACI, Brown Norway (BN), Buffalo (BUF), Fischer (FSH) and Wistar Kyoto (WKY) rats (the other three strains being no longer commercially available). We first tested the animals on the runway task, and found consistent differences between strains **(figure 5A)**. Most notably, BUF and BN rats had particularly high and low run latencies, averaging 586 ± 70 and 43 ± 13 seconds, respectively (n=8-12 per group, t(12)= 6.114, p<0.0001) **(figure 5B)**. From all five inbred strains, we used established formulas to estimate the within and between strain variances, deriving an estimated heritability of h^2^ = 0.28 [24]. As with the breeding experiment, this method showed that cocaine avoidance is heritable, albeit with the limitation that of only 5 of the 8 constituent HS strains were commercially available for us to test. We also tested the inbred strains on control tasks that could confound run latencies. BN and BUF rats were not different in either novelty induced locomotion (n=8-10 per group, t(16)=0.8241, p=0.4), or progressive punishment breakpoint, (n=6-8 per group, t(12)=0.7954, p=0.4). However, BUF rats had a higher progressive ratio breakpoint compared to the BN, suggesting that their higher run latencies are not due to decreased reward seeking (n=6-7 per group, t(11)=2.478, p=0.03) **(figure 5C)**. Altogether, these results again suggest that differences in runway latencies between the BN and the BUF are driven by the properties of cocaine, rather than generalized differences in locomotion, or processing of other rewards or costs.

**Figure 5.**
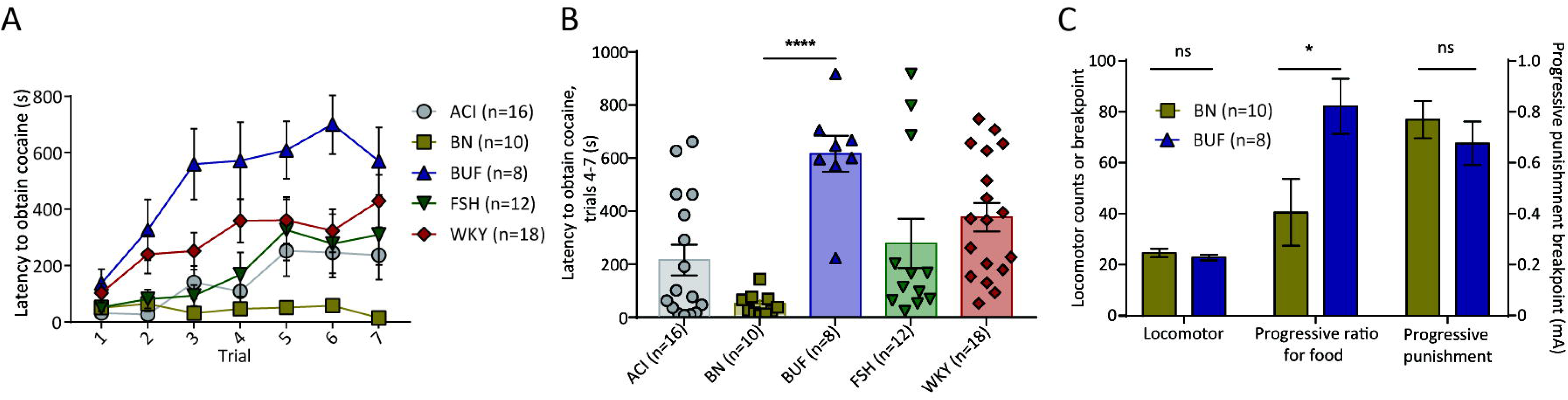
(**A**) We measured behavior on the runway task of five inbred strains: ACI, Brown Norway (BN), Buffalo (BUF), Fischer (FSH) and Wistar Kyoto (WKY), (**B**) Inbred groups had different average run latencies, with BUF and BN strains being the highest and lowest. (D) BN and BUF differed in their progressive ratio breakpoints, but not in their locomotor responses to novelty nor progressive punishment breakpoints. *\*p<0.05, ****p<0.0001.*

### Punishment resistance, another key contributor to addiction propensity, is heritable independently of cocaine avoidance or novelty-induced locomotion

Our results above examined punishment resistance as a control task to show that differences in runway behavior were not due to differences in generalized aversive learning. However, resistance to punishment is an important contributor to addiction in its own right, potentially also controlled by genetic factors. To assess the heritability of punishment resistance, we trained rats from all 5 inbred rat strains on the progressive punishment task described above. We found large strain differences in shock breakpoints that were not correlated with cocaine avoidance (n=6-7 per group, r^2^ = 0.031, p=0.78) **(figure 6A)** or locomotor response to novelty (n=6-7 per group, r^2^=0.032, p=0.77 **(figure 6B)**. For example, the Wistar-Kyoto (WKY) strain had a footshock breakpoint nearly quadruple those of the other strains (one-way ANOVA, Bonferroni test, n=6-10 per group, p<0.0001) **(figure 6C)**, even though they were not particularly extreme for run latency measures. From these results, we estimated a heritability of h^2^ = 0.62 for the punishment resistance phenotype [24]. Because punishment resistance is a general predisposing trait for all addictions, independently of any particular drug reinforcer, we had chosen to use a food rather than drug reward. Nevertheless, we still found that in WKY and ACI rats, shock breakpoints for cocaine paralleled those for food where the WKY rats had a shock breakpoint nearly twice as high as that of the ACI for cocaine reward (n=5-6 per group, t(9)=5.794, p=0.0003) **(figure 6D)**, suggesting that resistance to punishment varies independently of whether the reward is food or cocaine. Although we had hypothesized that the higher shock breakpoints indicate greater resistance to punishment, they are also consistent with an increased motivation to obtain food rewards. To tease apart these two possibilities, we tested all inbred strains on the progressive ratio (PR) food task mentioned above, and found that PR breakpoints did not correlate with shock breakpoints in the punishment resistance task (n=6 per group, r^2^=0.013, p=0.86) **(figure 6D)**, where WKY had a similar PR breakpoint as other strains. Finally, it remains possible that the WKY rats’ higher footshock breakpoints were simply due to deficiencies in pain perception. To assess pain sensitivity, we tested inbred animals on their latency to escape acute shocks of various intensities in a two-way shuttlebox, and found no differences between any of the 5 strains at any shock intensity (two-way ANOVA, Bonferroni test, n=5-6 per group, p>0.05), **(figure 6E)**.

**Figure 6.**
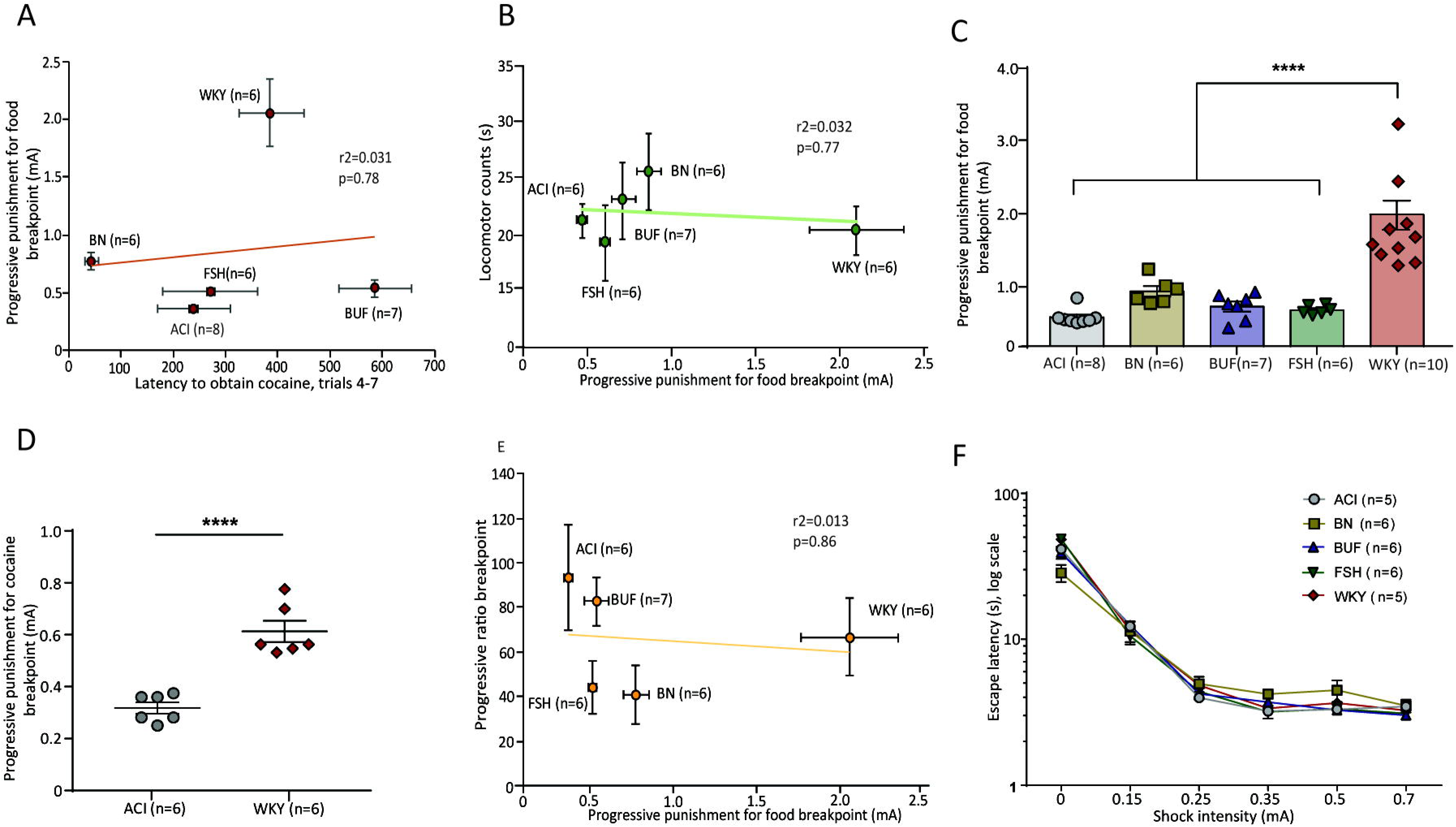
(**A**) Punishment resistance (for food) varied markedly between inbred strains, and varied independently of cocaine avoidance and (**B**) locomotor responses to novelty. (**C**) Shock break-point for food reward was significantly higher in WKY rats compared to the other four inbred strains. (**D**) WKY rats had significantly higher progressive punishment breakpoints for cocaine compared to ACI rats. (**E**) Progressive ratio breakpoints for food varied independently of progressive punishment breakpoint (**F**) Latencies to escape footshock did not differ between any inbred strains at any shock intensity. *\*\*\*\*p<0.0001.*

### High cocaine-avoiders have reduced rates of acquisition of cocaine self-administration but higher rates of reinstatement

After having extensively characterized cocaine avoidance, we next tested whether this phenotype predicts animals’ behavior in a conventional self-administration paradigm. We tested a new group of SD rats on the runway task then trained them to leverpress for cocaine in a self-administration paradigm, and found that run latencies were negatively correlated with lever pressing during the first three days of self-administration (n=22, r^2^=0.33, p=0.005, bar graph: t(20)=3.711, p=0.0014) **(Figure 7A)**. However, all animals eventually met the acquisition criteria, and the mean number of cocaine infusions for the last 3 days of self-administration was not significantly different between high- and low-avoiders (n=11-13 per group, t(22)=1.019, p=0.31) **(Figure 7B)**. We subsequently extinguished cocaine seeking behavior, and tested animals on both cued-induced and drug-primed reinstatement. Surprisingly, high avoiders were more likely to reinstate the extinguished cocaine seeking behavior compared to the low avoider group for both types of reinstatement (cued reinstatement: n=16, r^2^=0.48, p=0.003, bar graph: t(14)=2.659, p=0.019, cocaine-primed reinstatement: n=8, r^2^=0.60, p=0.02) **(Figure 7C and 7D)**.

**Figure 7.**
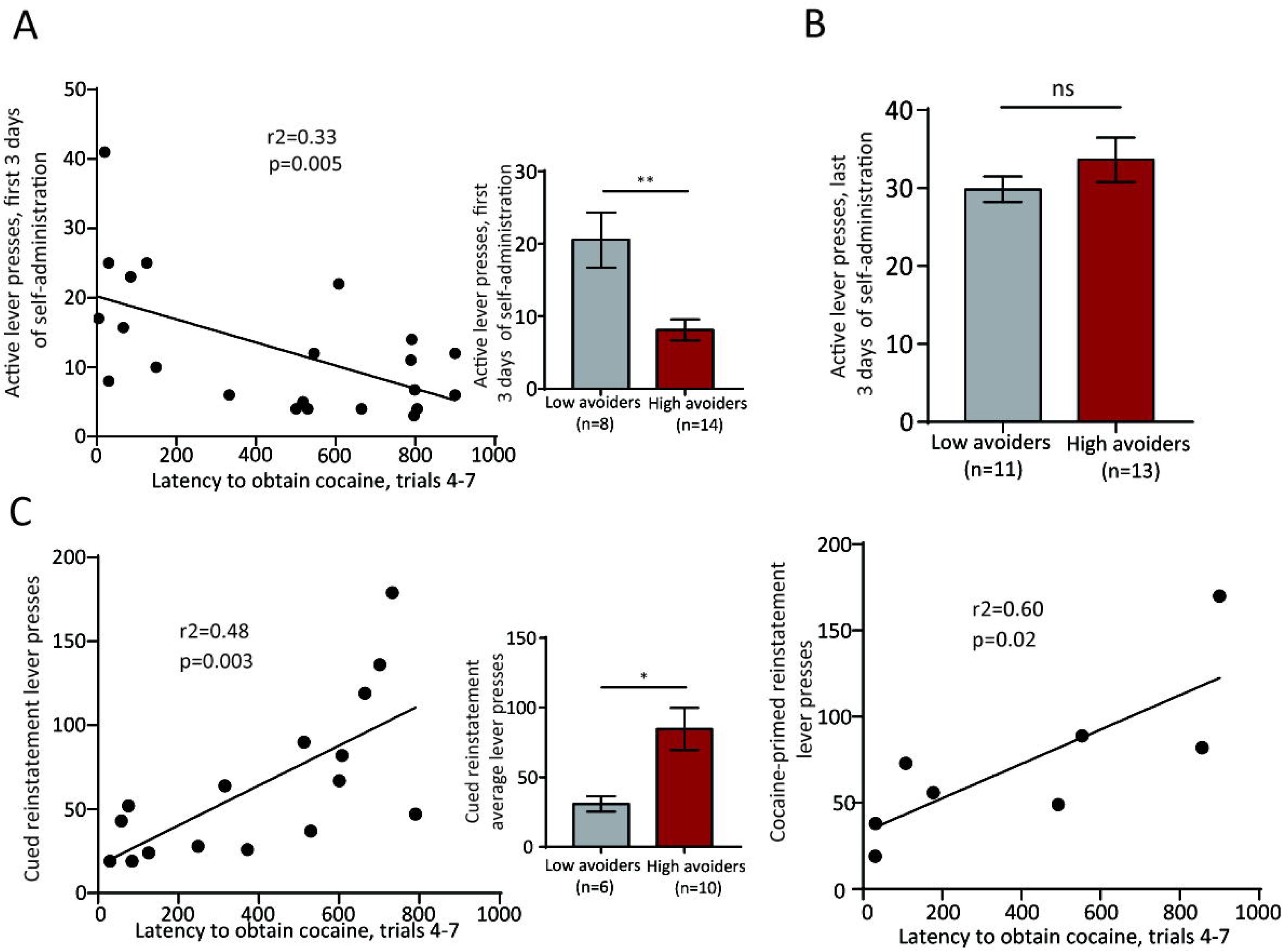
(**A**) Latencies to obtain cocaine (average of trials 4-7) correlated negatively with acquisition of lever pressing during the first three days of cocaine self-administration (**B**) The average number of cocaine infusions during the last 3 days of self-administration did not differ between high and low avoiders. Runway latencies to obtain cocaine correlated positively with both cued (**C**) and cocaine-primed (**D**) reinstatement lever pressing. *\*p<0.05, **p<0.01.*

## DISCUSSION

In this study, we found that conditioned avoidance to cocaine varies greatly between individual rats in both SD and HS strains, as measured using either a runway operant task or CPA. In both strains, we further found that cocaine avoidance responds rapidly to selective breeding, with large differences in offspring appearing after a single generation of selection. Together with additional experiments showing differences between several inbred rat strains, these data indicate that cocaine avoidance is strongly heritable. We also tested animals’ resistance to punishment and found it to also be highly heritable and to inherit independently of cocaine avoidance despite both behaviors being mediated by the RMTg [11, 20]. Indeed, we found that cocaine avoidance correlates strongly with cocaine-induced c-Fos induction in the RMTg, suggesting that individual variations in RMTg activation by cocaine might underlie individual differences in cocaine avoidance. Lastly, we showed that individuals showing high levels of avoidance of cocaine were slower to acquire intravenous cocaine self-administration in a standard operant leverpress cocaine-seeking task but reinstated more strongly to both cues and cocaine priming infusions, suggesting that cocaine’s aversive effects have opposite influences on two distinct stages of drug-seeking behavior.

Taken together, these data suggest that the transition from low to high rates of drug use may occur in multiple stages, and that these stages may be independently governed by distinct brain mechanisms controlled by distinct, albeit unidentified, genes. These results parallel well-known phenomena in humans, as many humans stop using drugs on their own prior to addiction onset, while others progress onto heavy use and dependence. Furthermore, epidemiological studies suggest that each of the important stages of addiction, including vulnerability to initiation, continued drug-seeking, and propensity to become dependent, are under some genetic influence, where the genetic etiology becomes progressively more important as the individual progresses from first use to dependence [25-34]. Indeed, family, adoption and twin studies of illicit drug use have calculated heritability estimates ranging from 30% to 80%, with cocaine addiction having particularly high heritability at 72% [35-38]. Although genetic studies have identified several common variants associated with substance abuse [39-43], the field has been characterized by lack of replication and there are remarkably few genes that can be firmly associated with addiction. Indeed, studying the genetics of complex disorders, such as addiction, in a human population poses several challenges [35], as addiction is an intricate disorder with numerous interacting facets, including environmental factors, drug-induced neurobiological changes, comorbidity with other neuropsychiatric disorders and personality traits [44-48]. Multiple genetic variants that affect any of these factors may work in synergy to affect vulnerability and severity of addiction. Hence, we sought strategies to examine addiction heritability in an animal model in which the complexity of the addictive phenotype can be subdivided into smaller constituents that could be investigated independently, using tasks that deconstruct the cost-benefit decision processes into its building blocks.

We measured one constituent part of addiction vulnerability using the runway task of cocaine seeking. Although it is not as widely used as lever pressing for cocaine self-administration, we chose it for several reasons. First, the single-trial nature of the task isolates the aversive effects of cocaine, ensuring that they are not masked by subsequent doses as would happen in lever pressing paradigms. Second, the limited availability of cocaine more closely resembles earlier stages of drug use where trial doses are highly intermittent, unlike the unbounded amounts available in conventional self-administration paradigms. Despite these differences, we also found that self-administration using the runway task and traditional lever-based self-administration are correlated, demonstrating that both paradigms share some of the same underlying processes.

While conditioned avoidance to cocaine may strongly influence early stages of drug-seeking, after drug acquisition, animals are able to mitigate the crash with subsequent doses. Hence, continuation of drug-seeking in later stages of addiction may depend on the degree to which negative consequences of drug-seeking are able to curtail this behavior. Indeed, a defining feature of an addictive behavior is its persistence despite negative financial, social and medical consequences. This characteristic applies to both drug and non-drug addictions [49, 50], such as gambling addictions, which are also strongly heritable (h^2^ = 0.49) [51-53]. Because this failure to learn from punishment is shared between non-drug and drug addictions, as well as underlying neural circuitry elements [50, 54], a key element of this study was the finding of strong heritability of punishment learning. Notably, cocaine avoidance and punishment resistance appear to inherit independently, as there is no correlation between the two, either across strains or individual rats. This independence occurs despite both tasks depending on the RMTg, a brain region carries an aversive “teaching signal” driving avoidance learning, including avoidance of cocaine and the ability to withhold food-seeking in the face of a footshock punisher [11, 14, 55]. The current study further found evidence that cocaine activates the RMTg more strongly in rats that show avoidance to the drug. Together with prior studies showing that RMTg disruption entirely abolishes conditioned avoidance of cocaine, these results suggest individual variation in RMTg activation might underlie individual variation in cocaine avoidance behavior (though perhaps not punishment resistance).

Finally, we showed that the runway model has important clinical significance in terms of predicting drug acquisition and reinstatement. Indeed, high-avoiders initially did not acquire self-administration and were thus slower to transition to regular use, suggesting that cocaine aversion may serve as a protective factor against addiction. However, avoidance behavior correlated with reinstatement suggesting that, as has been previously hypothesized, addictive behavior is partly driven by negative reinforcement that emerges from an individual’s attempt to reduce aversive motivational states arising in the absence of the drug [56, 57].

Our data have given us clues to the anatomical substrates underlying avoidance learning and addiction, but further study is needed to identify the specific molecular targets within these circuits that behave so differently across individuals, thereby improving our understanding of the biochemical processes underlying addiction vulnerability, while also improving our ability to predict individual vulnerabilities to risk and to recovery.

## Materials and methods

### Animals

Adult male Sprague Dawley, Heterogeneous Stock, and inbred animals weighing 250–300g upon delivery from vendor were individually housed in standard shoebox cages with food and water provided *ad libitum*, unless otherwise stated. Procedures conformed to the National Institutes of Health *Guide for the Care and Use of Laboratory Animals*, and all protocols were approved by Medical University of South Carolina Institutional Animal Care and Use Committee.

### Surgery

Rats were anesthetized with isoflurane and fitted with an indwelling intravenous catheter inserted into the right jugular vein, and subcutaneously passed to a guide cannula that exited on the animal’s back. After surgery, catheter patency was maintained through daily flushing with 0.05 ml of TCS. Animals recovered for 7 days prior to behavioral testing. Catheter patency was assessed periodically through observation of the loss of the righting reflex after i.v. injection of, methohexital (Brevital, 2.0 mg/kg/0.1 ml). Rats that were unresponsive to Brevital were implanted with a new catheter using the left jugular vein.

### Behavioral testing

#### Runway operant cocaine-seeking

Methods are similar to our previous work [11]. Briefly, the runway consists of two opaque plastic compartments (“start” and “goal”) connected by a 170cm long corridor. Rats are tethered to an intravenous line, placed into the start box, after which doors are opened, allowing free exploration of the apparatus. Entry into the goal box causes doors to close and a syringe pump to deliver cocaine (0.75mg/kg iv, same dose as in CPP/CPA). Failure to enter the goal compartment after 15 minutes resulted in a timeout. 7 trials were given at least 4 hours apart, ensuring that animals ran in a cocaine-free state.

#### Progressive ratio (PR) and punishment resistance

Training procedures are identical to those previously described by our lab [14]. Briefly, animals are food restricted to 85% of initial body weight, and then trained to leverpress for food pellets (45mg). After stable response rates are reached, animals are tested with either a progressively increasing effort requirement (i.e. more leverpresses required to obtain each pellet) or with progressively increasing punishment (i.e. a footshock of increasing intensity presented just after food pellet delivery). Both the effort requirement and shock intensity increase on a trial-by-trial basis as previously described [58]. If rats time out before pressing a failure is noted. For both tasks, the breakpoint is the last successfully completed ratio or shock intensity endured before timeout.

#### Locomotor

Locomotor activity was recorded in sound-attenuating cabinets where the operant chambers were housed. Each chamber was equipped with an array of 4 photodetector-emitter pairs. Movement within the chamber produced photobeam interruptions that were recorded. Recordings lasted for 30 minutes and the average of two sessions were collected on two different days.

#### CPP/CPA

On day 1, animals were placed in the central chamber and a baseline was collected for 15 min. Animals received a total of four conditioning sessions for each task. Rats received an infusion of i.v. cocaine (0.75 mg/kg) on the paired side (0 min post-infusion for CPP and 15 min post-infusion for CPA) and saline on the unpaired trials (left for 10 min in the chamber). Two session were conducted daily (cocaine and saline) counterbalanced in order. In our group of animals, the chambers were counterbalanced as well. The day after the final conditioning session (i.e., test day: day 5), animals were placed into the central compartment of the CPP apparatus for 15 min, and time spent in each compartment was recorded.

#### Shuttlebox shock escape

Rats are placed into a shuttlebox having two equal sized compartments and a motorized doorway connecting the two. At two-minute intervals, footshock is delivered to the rat, which is terminated after the rat crosses through the doorway. Photobeams measure the latency to reach the other side. 10 trials are given for each session, and average latency is recorded for each rat. The shock intensity is fixed during each session, and two sessions are conducted for each rat at each shock intensity to reduce session-by-session variability. Different shock intensities were used (0mA, i.e. control, 0.15mA, 0.25mA, 0.35Ma, 0.5mA and 0.7mA). Shocks are tested in ascending order, to avoid carryover of fear conditioning from one session to the next.

#### Self-administration

The task is performed in operant chambers equipped with two levers, a house light, cue light, and tone generator. The designated active lever delivered an infusion of cocaine (0.75 mg/kg/infusion) along with a compound cue (light + tone), followed by a 20 s time out. Before going through extinction, animals must have had at least 10 days of self-administration with at least 10 infusions. After self-administration, rats were extinguished for at least 10 days until criterion was reached (2 days <10 presses on the active lever) and given cue induced or drug-primed reinstatement tests.

### Histology

Rats were transcardially perfused with saline and 10% formalin. Brains were removed and placed in 10% formalin overnight before storage in 20% sucrose with PBS azide. c-Fos counts were blindly performed immunohistochemically for staining of cocaine (10mg/kg; i.p.)-induced rabbit anti-c-Fos (1:10K, EMD Millipore, MA, USA).

## References

1. Wenzel, J.M., et al., Effects of lidocaine-induced inactivation of the bed nucleus of the stria terminalis, the central or the basolateral nucleus of the amygdala on the opponent-process actions of self-administered cocaine in rats. Psychopharmacology (Berl), 2011. 217(2): p. 221–30.

2. Heyman, G.M., Addiction and choice: theory and new data. Front Psychiatry, 2013. 4: p. 31.

3. Anthony, J.C., L.A. Warner, and R.C. Kessler, Comparative epidemiology of dependence on tobacco, alcohol, controlled substances, and inhalants: Basic findings from the National Comorbidity Survey. Experimental and Clinical Psychopharmacology, 1994. 2(3): p. 244–268.

4. Goldman, D., G. Oroszi, and F. Ducci, The genetics of addictions: uncovering the genes. Nat Rev Genet, 2005. 6(7): p. 521–32.

5. Ho, M.K., et al., Breaking barriers in the genomics and pharmacogenetics of drug addiction. Clin Pharmacol Ther, 2010. 88(6): p. 779–91.

6. Ducci, F. and D. Goldman, The genetic basis of addictive disorders. Psychiatr Clin North Am, 2012. 35(2): p. 495–519.

7. Stead, J.D., et al., Selective breeding for divergence in novelty-seeking traits: heritability and enrichment in spontaneous anxiety-related behaviors. Behav Genet, 2006. 36(5): p. 697–712.

8. Ettenberg, A., et al., Evidence for opponent-process actions of intravenous cocaine. Pharmacol Biochem Behav, 1999. 64(3): p. 507–12.

9. Ettenberg, A., et al., On the positive and negative affective responses to cocaine and their relation to drug self-administration in rats. Psychopharmacology (Berl), 2015. 232(13): p. 2363–75.

10. Geist, T.D. and A. Ettenberg, Concurrent positive and negative goalbox events produce runway behaviors comparable to those of cocaine-reinforced rats. Pharmacol Biochem Behav, 1997. 57(1-2): p. 145–50.

11. Jhou, T.C., et al., Cocaine drives aversive conditioning via delayed activation of dopamine-responsive habenular and midbrain pathways. J Neurosci, 2013. 33(17): p. 7501–12.

12. Maroteaux, M. and M. Mameli, Cocaine evokes projection-specific synaptic plasticity of lateral habenula neurons. J Neurosci, 2012. 32(36): p. 12641–6.

13. Ettenberg, A., Opponent process properties of self-administered cocaine. Neurosci Biobehav Rev, 2004. 27(8): p. 721–8.

14. Vento, P.J., et al., Learning from one’s mistakes: A dual role for the rostromedial tegmental nucleus in the encoding and expression of punished reward seeking. Biol Psychiatry, 2016.

15. Li, H., et al., Generality and opponency of rostromedial tegmental (RMTg) roles in valence processing. Elife, 2019. 8.

16. Jhou, T.C., et al., The rostromedial tegmental nucleus (RMTg), a GABAergic afferent to midbrain dopamine neurons, encodes aversive stimuli and inhibits motor responses. Neuron, 2009. 61(5): p. 786–800.

17. Jhou, T.C., et al., The mesopontine rostromedial tegmental nucleus: A structure targeted by the lateral habenula that projects to the ventral tegmental area of Tsai and substantia nigra compacta. J Comp Neurol, 2009. 513(6): p. 566–96.

18. Lopez-Quintero, C., et al., Probability and predictors of remission from life-time nicotine, alcohol, cannabis or cocaine dependence: results from the National Epidemiologic Survey on Alcohol and Related Conditions. Addiction, 2011. 106(3): p. 657–69.

19. Klingemann, H.K., The motivation for change from problem alcohol and heroin use. Br J Addict, 1991. 86(6): p. 727–44.

20. Vento, P.J., et al., Learning From One’s Mistakes: A Dual Role for the Rostromedial Tegmental Nucleus in the Encoding and Expression of Punished Reward Seeking. Biol Psychiatry, 2017. 81(12): p. 1041–1049.

21. Woolverton, W.L. and C.R. Schuster, Intragastric self-administration in rhesus monkeys under limited access conditions: methodological studies. J Pharmacol Methods, 1983. 10(2): p. 93–106.

22. Solberg Woods, L.C., et al., Fine-mapping a locus for glucose tolerance using heterogeneous stock rats. Physiol Genomics, 2010. 41(1): p. 102–8.

23. Solberg Woods, L.C., et al., Heterogeneous stock rats: a new model to study the genetics of renal phenotypes. Am J Physiol Renal Physiol, 2010. 298(6): p. F1484–91.

24. Hegmann, J.P. and B. Possidente, Estimating genetic correlations from inbred strains. Behav Genet, 1981. 11(2): p. 103–14.

25. McGue, M., R.W. Pickens, and D.S. Svikis, Sex and age effects on the inheritance of alcohol problems: a twin study. J Abnorm Psychol, 1992. 101(1): p. 3–17.

26. True, W.R., et al., Genetic and environmental contributions to smoking. Addiction, 1997. 92(10): p. 1277–87.

27. Juli, G. and L. Juli, Genetic of addiction: common and uncommon factors. Psychiatr Danub, 2015. 27 Suppl 1: p. S383–90.

28. Kendler, K.S., et al., Genetic and environmental influences on alcohol, caffeine, cannabis, and nicotine use from early adolescence to middle adulthood. Arch Gen Psychiatry, 2008. 65(6): p. 674–82.

29. Johnson, E.O., M.B. van den Bree, and R.W. Pickens, Indicators of genetic and environmental influence in alcohol-dependent individuals. Alcohol Clin Exp Res, 1996. 20(1): p. 67–74.

30. Kendler, K.S., et al., A population-based twin study in women of smoking initiation and nicotine dependence. Psychol Med, 1999. 29(2): p. 299–308.

31. Heath, A.C., et al., Estimating two-stage models for genetic influences on alcohol, tobacco or drug use initiation and dependence vulnerability in twin and family data. Twin Res, 2002. 5(2): p. 113–24.

32. Neale, M.C., et al., Extensions to the modeling of initiation and progression: applications to substance use and abuse. Behav Genet, 2006. 36(4): p. 507–24.

33. Agrawal, A., et al., Illicit drug use and abuse/dependence: modeling of two-stage variables using the CCC approach. Addict Behav, 2005. 30(5): p. 1043–8.

34. Merikangas, K.R. and S. Avenevoli, Implications of genetic epidemiology for the prevention of substance use disorders. Addict Behav, 2000. 25(6): p. 807–20.

35. Agrawal, A., et al., The genetics of addiction-a translational perspective. Transl Psychiatry, 2012. 2: p. e140.

36. Kendler, K.S., et al., Illicit psychoactive substance use, heavy use, abuse, and dependence in a US population-based sample of male twins. Arch Gen Psychiatry, 2000. 57(3): p. 261–9.

37. Tsuang, M.T., et al., The Harvard Twin Study of Substance Abuse: what we have learned. Harv Rev Psychiatry, 2001. 9(6): p. 267–79.

38. van den Bree, M.B., et al., Genetic and environmental influences on drug use and abuse/dependence in male and female twins. Drug Alcohol Depend, 1998. 52(3): p. 231–41.

39. Edenberg, H.J., The genetics of alcohol metabolism: role of alcohol dehydrogenase and aldehyde dehydrogenase variants. Alcohol Res Health, 2007. 30(1): p. 5–13.

40. Thorgeirsson, T.E., et al., Sequence variants at CHRNB3-CHRNA6 and CYP2A6 affect smoking behavior. Nat Genet, 2010. 42(5): p. 448–53.

41. Zhang, H., et al., Pro-opiomelanocortin gene variation related to alcohol or drug dependence: evidence and replications across family- and population-based studies. Biol Psychiatry, 2009. 66(2): p. 128–36.

42. Kalayasiri, R., et al., Dopamine beta-hydroxylase gene (DbetaH) -1021C-->T influences self-reported paranoia during cocaine self-administration. Biol Psychiatry, 2007. 61(11): p. 1310–3.

43. Uhl, G.R., I. Sora, and Z. Wang, The mu opiate receptor as a candidate gene for pain: polymorphisms, variations in expression, nociception, and opiate responses. Proc Natl Acad Sci U S A, 1999. 96(14): p. 7752–5.

44. Stone, A.L., et al., Review of risk and protective factors of substance use and problem use in emerging adulthood. Addict Behav, 2012. 37(7): p. 747–75.

45. Wong, C.C. and G. Schumann, Review. Genetics of addictions: strategies for addressing heterogeneity and polygenicity of substance use disorders. Philos Trans R Soc Lond B Biol Sci, 2008. 363(1507): p. 3213–22.

46. Bau, C.H., et al., Heterogeneity in early onset alcoholism suggests a third group of alcoholics. Alcohol, 2001. 23(1): p. 9–13.

47. Belin, D., et al., In search of predictive endophenotypes in addiction: insights from preclinical research. Genes Brain Behav, 2016. 15(1): p. 74–88.

48. Gottesman, II and T.D. Gould, The endophenotype concept in psychiatry: etymology and strategic intentions. Am J Psychiatry, 2003. 160(4): p. 636–45.

49. Potenza, M.N., Review. The neurobiology of pathological gambling and drug addiction: an overview and new findings. Philos Trans R Soc Lond B Biol Sci, 2008. 363(1507): p. 3181–9.

50. Potenza, M.N., Should addictive disorders include non-substance-related conditions? Addiction, 2006. 101 Suppl 1: p. 142–51.

51. Slutske, W.S., et al., Genetic and environmental influences on disordered gambling in men and women. Arch Gen Psychiatry, 2010. 67(6): p. 624–30.

52. Slutske, W.S., et al., Common genetic vulnerability for pathological gambling and alcohol dependence in men. Arch Gen Psychiatry, 2000. 57(7): p. 666–73.

53. Eisen, S.A., et al., The genetics of pathological gambling. Semin Clin Neuropsychiatry, 2001. 6(3): p. 195–204.

54. Leeman, R.F. and M.N. Potenza, Similarities and differences between pathological gambling and substance use disorders: a focus on impulsivity and compulsivity. Psychopharmacology (Berl), 2012. 219(2): p. 469–90.

55. Wenzel, J.M., et al., Effects of lidocaine-induced inactivation of the bed nucleus of the stria terminalis, the central or the basolateral nucleus of the amygdala on the opponent-process actions of self-administered cocaine in rats. Psychopharmacology (Berl), 2011.

56. Koob, G.F., et al., Addiction as a stress surfeit disorder. Neuropharmacology, 2014. 76 Pt B: p. 370–82.

57. Wheeler, D.S., et al., Drug predictive cues activate aversion-sensitive striatal neurons that encode drug seeking. J Neurosci, 2015. 35(18): p. 7215–25.

58. Richardson, N.R. and D.C. Roberts, Progressive ratio schedules in drug self-administration studies in rats: a method to evaluate reinforcing efficacy. J Neurosci Methods, 1996. 66(1): p. 1–11.

